# Selective inhibition of excitatory synaptic transmission alters the emergent bursting dynamics of in vitro neural networks

**DOI:** 10.1101/2022.09.08.507095

**Authors:** Janelle Shari Weir, Nicholas Christiansen, Axel Sandvig, Ioanna Sandvig

## Abstract

Neurons in vitro connect to each other and form neural networks that display emergent electrophysiological activity. This activity begins as spontaneous uncorrelated firing in the early phase of development, and as functional excitatory and inhibitory synapses mature, the activity typically emerges as spontaneous network bursts. Network bursts are events of coordinated global activation among many neurons interspersed with periods of silencing and are important for synaptic plasticity, neural information processing, and network computation. While bursting is the consequence of balanced excitatory-inhibitory (E/I) interactions, the functional mechanisms underlying their evolution from physiological to potentially pathophysiological states, such as decreasing or increasing in synchrony, are still poorly understood. Synaptic activity, especially that related to maturity of E/I synaptic transmission, is known to strongly influence these processes. In this study, we used selective chemogenetic inhibition to target and disrupt excitatory synaptic transmission in in vitro neural networks to study functional response and recovery of spontaneous network bursts over time. We found that over time, inhibition resulted in increases in both network burstiness and synchrony. Our results indicate that the disruption in excitatory synaptic transmission during early network development likely affected inhibitory synaptic maturity which resulted in an overall decrease in network inhibition at later stages. These findings lend support to the importance of E/I balance in maintaining physiological bursting dynamics and, conceivably, information processing capacity in neural networks.

## 1. Introduction

Neural network dynamics emerge over the course of development in vitro. Spontaneous network activity starts as immature tonic spiking and primitive patterns of synchronized activity in the early phases of development (Ben-Ari, 2001) which then progresses towards more complex behavior characterized by bursts (van Pelt et al., 2004, Fardet et al., 2018). Typically, in vitro neural networks start exhibiting bursts between 6 and 14 DIV (Chiappalone et al., 2006, Wagenaar et al., 2006). Such early bursts, described as “superbursts” (Stephens et al., 2012), are posited to be driven by depolarizing gamma – aminobutyric acid type A (GABA_A_) receptors and are hallmarks of early network development. At this stage, neuronal interactions are strengthened leading to recurrent coactivation among several neurons, which manifest as network bursts. These network bursts become more recurring as the neural network reaches maturity around 21 DIV and onwards, with burst profile of higher frequency, shorter burst onset and offset, and shorter duration (Bisio et al., 2014, Chiappalone et al., 2006).

Network bursts are shown to be driven by excitatory synaptic transmission (Robinson et al., 1993, Kudela et al., 2003, Teppola et al., 2019), primarily mediated by glutamatergic ionotropic N-methyl-D-aspartate (NMDA) receptors and alpha-amino-3-hydroxy-5-methyl-4-isoxazolepropionic acid (AMPA) receptors. Fast inhibition by GABA_A_ receptors also mediates network activity and burst emergence by maintaining a balance in excitatory – inhibitory (E/I) synaptic transmission (Teppola et al., 2019). Early in vitro studies reported that the relationship between network age, structure and the resulting activity is due to variations in synaptic connections and the differential developmental periods of excitatory and inhibitory synaptic transmission (Burgard and Hablitz, 1994, Kamioka et al., 1996). As the network achieves adequate interconnectivity and inhibitory synapses become more functionally mature during the later stages of development, network dynamics are reported to progress from spontaneous uncorrelated firing to more complex patterns of synchronized network bursts (Kamioka et al., 1996, Opitz et al., 2002, Wagenaar et al., 2006, Baltz et al., 2010). It has been suggested that the propagation of synchronized bursts plays an important role in shifting the network from immaturity into a stage characterized by a highly diversified range of electrical signaling (Ben-Ari, 2001), rendering the network capable of complex information processing and encoding. Several in vivo studies have reported similar age specific correlation of the emergence of network bursts with functional circuit development in various parts of the nervous system including the hippocampus (Blankenship and Feller, 2010, Raus Balind et al., 2019), cerebellar cortex (Dizon and Khodakhah, 2011, Hoehne et al., 2020), visual cortex (Chiu and Weliky, 2001), medulla (Pena and Ramirez, 2004, Magalhaes et al., 2021) and spinal cord (Darbon et al., 2004). These findings suggest that excitatory and inhibitory synaptic maturity are important drivers of network bursts, burst characteristics and subsequent network function. The effect of selective disruption of E/I balance on bursting dynamics in neural networks may therefore reveal substantial biological insights into network function, adaptability, and robustness.

Investigating inhibitory – excitatory synaptic contribution to network burst evolution in vivo is challenging. This is in part because the brain comprises numerous complex multi-layered neural networks, with heterogeneous synaptic connectivity among subsets of burst-generating neurons that contribute to the dynamics of the network (Zeldenrust et al., 2018). The interweaving of different neurons and synapses at various topological and temporal scales makes it challenging to determine the relative impact of synaptic activity on physiological and pathophysiological bursting activity. Since in vitro neural networks represent a monolayered reductionist model of a brain network – while still maintaining salient age dependent electrophysiological dynamics (Ben-Ari, 2001, Chiappalone et al., 2006, Chiappalone et al., 2007, Sun et al., 2010, Schroeter et al., 2015) – the complexity is reduced along spatial dimensions, and thus enables study and selective manipulation in a controlled manner (Marom and Shahaf, 2002). Many studies have taken advantage of such reductionist in vitro models to investigate network burst dynamics at the synaptic level via manipulation that changes the balance between excitatory and inhibitory synaptic transmission. Methods such as pharmacological blockade of NMDA and AMPA receptors (Chub and O’Donovan, 1998, Li et al., 2007, Suresh et al., 2016) and membrane current blockers (Ramakers et al., 1990, Ramakers et al., 1994, van Drongelen et al., 2006) have provided significant insights into the functional contribution of synaptic receptors and intrinsic membrane currents to the generation, maintenance, duration, and propagation of network bursts. In this study, we utilized hM4Di designer receptors exclusively activated by designer drugs (DREADDs) (Armbruster et al., 2007, Alexander et al., 2009, Urban and Roth, 2015, Khambhati and Bassett, 2016, Whissell et al., 2016, Panthi and Leitch, 2019, Haaranen et al., 2020a, Haaranen et al., 2020b, Lebonville et al., 2020, Ozawa and Arakawa, 2021) to selectively inhibit excitatory synaptic transmission – via G-protein coupled receptors (GPCRs) in calcium/calmodulin-dependent protein kinase alpha (CaMKlla) expressing neurons – in neural networks interfaced with microelectrode arrays (MEAs). Networks were chemogenetically inhibited at 14 DIV, 21 DIV and 28 DIV and their dynamics characterized in comparison to their baseline activity and to PBS vehicle and control, unperturbed networks. The internal characteristics of network bursts both during treatment (functional response to perturbation) and post treatment (recovery of the network) were analyzed. We found that inhibition of excitatory synaptic transmission increased bursting activity, as well as increased network synchronization within the chemogenetically inhibited networks by 28 DIV. Our results suggest that the long-term maintenance of the E/I balance depends on ongoing excitatory synaptic activity, and that disruption impairs physiological processes involved in modulating synchrony in maturing neural networks.

## 2. Materials and Methods

### 2.1 Culture of cortical networks on microelectrode arrays

Primary rat (Sprague Dawley) cortex neurons were obtained from ThermoFisher Scientific, US (Cat. No: A36511). Cells were thawed and seeded as a co-culture with 15% rat primary cortical astrocytes also from ThermoFisher Scientific (Cat. No: N7745100). The cells were plated at a density of approximately 1000 cells/mm^2^ on Nunc™ Lab -Tek™ chamber slides (Cat. No. 177380) coated with Geltrex matrix (cat. No. A1413201) at a working concentration of 0.5ug/cm^2^ for 1:100 dilution, both obtained from ThermoFisher Scientific. Pre-sterilized 6-well CytoView MEA plates were purchased from Axion Biosystems and coated with 0.5% polyethyleneimine diluted in HEPES (both from Sigma-Aldrich) and 20 μg/mL natural mouse laminin (ThermoFisher Scientific) diluted in DPBS according to the Axion coating protocol (Axion BioSystems, Georgia, USA). Cells were plated directly over the electrodes on Axion MEA plates at a density of approximately 1500 cells/mm^2^ and incubated for 4 hours before topping up wells to 1mL with media. Cells were plated and maintained in Neurobasal™ Plus Medium supplemented with 2% B-27 Plus Supplement and 0.5% GlutaMAX™ all from ThermoFisher Scientific. The culture media was also supplemented with 0.2% (1:500 dilution from a 5 μg/ml working concentration) Plasmocin™ Prophylactic (ant. mpp; InvivoGen). The day of plating from cryopreservation was allocated as day 0 and 50% media changes were carried out every 2-3 days. Cells were always kept in a 5%CO_2_ incubator at 37°C except during media changes and imaging. All the wells on a single Axion MEA plate were allocated to one experimental condition. This ensured that networks that received the DREADDs virus were handled separately from the control networks, which did not receive the virus.

### 2.2 AAV 2/1 hM4Di CaMKlla-DREADDs production and in vitro transduction

Vector production and purification was performed in-house at the Viral Vector Core Facility (Kavli Institute, NTNU). Tittering of the viral stock was determined as approximately 10^11^ vg/mL. High viral stocks were aliquoted into 20ul volumes and stored at −80 °C. Aliquots for use were thawed on ice and remaining virus aliquoted at store at −80°C. The maximum number of thaws for any aliquot used was 3 times. At 7 DIV, the neurons were transduced by removing 80% of the cell media from the culture and directly adding a dilution of AAV viral particles encoding experimental hM4Di -CaMKlla-DREADDs to the neurons (**Figure 1A**). The titer of the viral dilution used for each well was 1 × 10^3^ viral units per neuron based on tests at different viral concentrations (results not included). The cultures were gently agitated for 30 seconds to ensure proper distribution of the viral particles and then incubated for 8 hours. Afterwards, each well was topped up to 1mL with fresh media without Plasmocin™ Prophylactic and incubated for an additional 40 hours in 5%CO_2_, 37°C incubator. After the incubation period, 50% media changes were carried out as scheduled. The vector encodes mCherry which is a bright red fluorescent protein tag that makes it possible to visualize results soon after transduction (**Figure 1B**).

**Figure 1.**
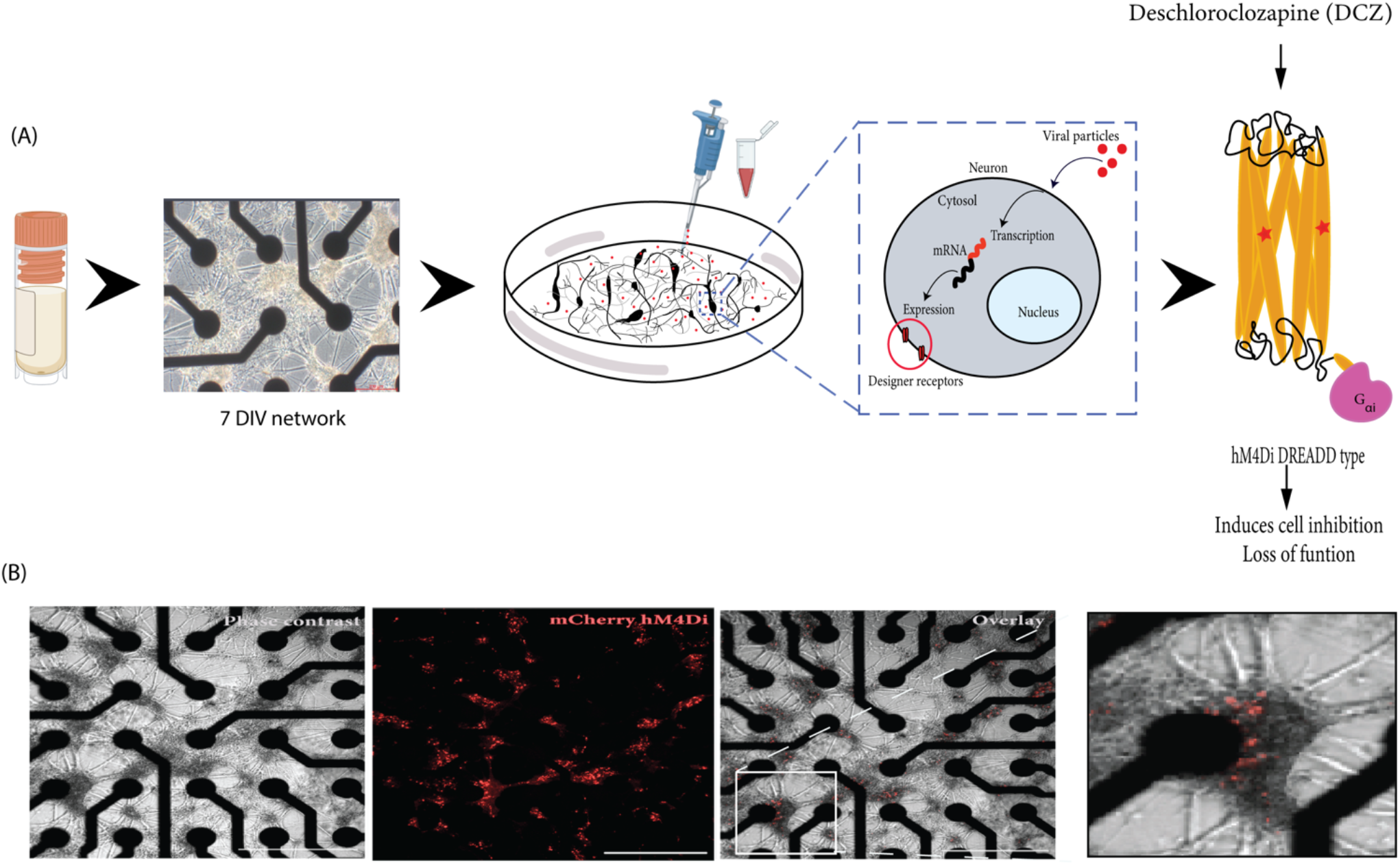
Illustration of the workflow of the study. **(A)** Cryopreserved cortical neurons were thawed and seeded on precoated MEAs until 7 DIV (scale bar = 200 μm). Neurons grow on top of the electrodes (black) and connect with each other across the surface. The DREADD protein is encoded in a replication deficient adeno-associated viral vector (AAV2/1) that is targeted for cytoplasmic gene delivery, thus circumventing genomic integration. The vector was added directly to the neurons in culture (on the MEA). The vector codes for the cell-specific promoter CaMKll so that mRNA transcription is targeted specifically at excitatory neurons which will be the cells that express the designer receptors. The designed DREADD hM4Di receptor has mutations at two points, which results in the receptor being insensitive to its endogenous agonist and neurotransmitter acetylcholine and instead respond only to a physiologically inert exogenous molecule (designer drug) such as deschloroclozapine (DCZ). When DCZ binds, the hM4DiR preferentially signals through the Gai/o subset of G-protein to inhibit adenylate cyclase and downstream cyclic adenosine monophosphate (cAMP) production, causing neuronal hyperpolarization and induces loss of cellular activity. **(B)** In vitro neural network on MEA at 12 days post adding the virus. DREADDs expression was confirmed without immunocytochemistry based on strong mCherry fluorescent expression in the network. Scale bar = 1250μm

### 2.3 Immunocytochemistry

At 14 DIV, parallel hM4Di DREADD networks were immunolabeled to investigate the specificity for vector mediated hM4Di expression in the CaMKlla positive neurons. The cultures were fixed with 4% Paraformaldehyde (PFA) for 20 minutes and washed with DPBS before cultures were permeabilized with a blocking solution of 0.03% Triton X-100 and 5% goat serum in DPBS for 2 hours at room temperature. Following blocking, antibodies at the indicated solutions **(Table 1)** were added in a buffer of 0.01% Triton X-100 and 1% goat serum in DPBS. Nuclei were stained with Hoechst (bisbenzimide H 33342 trihydrochloride, 14533, Sigma-Aldrich, 1: 5000 dilution). Samples were washed, mounted on glass cover slides with anti-fade fluorescence mounting medium (ab104135, Abcam) and imaged. All sample images were acquired using the EVOS M5000 imaging system (Invitrogen, ThermoFisher Scientific). Images were processed using Fiji/ImageJ and Adobe Illustrator 2020 version: 24.0.0.

**Table 1.** Overview of primary and secondary antibodies, species, and concentration.

| Primaries |  |  | Secondaries |  |  |
| --- | --- | --- | --- | --- | --- |
| Markers | Catalogue # | Concentration | Fluorescent | Catalogue # | Concentration |
| Ck mCherry | Ab205402<br>(Abcam) | 1:1000 | Goat-anti-Chicken<br>AlexaFluor 568 | Ab175477<br>(Abcam) | 1:1000 |
| Ms Calmodulin<br>(CaMKII) | MA3-918<br>(Invitrogen) | 1:250 | Goat-anti-Mouse<br>AlexaFluor 568 | A11019<br>(Invitrogen) | 1:1000 |
| Ms NMDAR1 | 32-0500<br>(Invitrogen) | 1:100 | Goat-anti-Mouse<br>AlexaFluor 647 | A21236<br>(Invitrogen) | 1:1000 |
| Ms GABA BR1 | Ab55051 (Abcam) | 1:250 | Goat-anti-Mouse<br>AlexaFluor 488 | A11001<br>(Invitrogen) | 1:1000 |
| Rb Calmodulin<br>(CaMKII) | Ab134041<br>(Abcam) | 1:200 | Goat-anti-Rabbit<br>AlexaFluor 568 | A11011<br>(Invitrogen) | 1:1000 |
| Rb Glutamate<br>decarboxylase<br>(GAD65/67) | Ab183999<br>(Abcam) | 1:100 | Goat-anti-Rabbit<br>AlexaFluor 647 | A21244<br>(Invitrogen) | 1:1000 |
| Rb Map2 | Ab32454 (Abcam) | 1:250 | Goat-anti-Rabbit<br>AlexaFluor 488 | A11008<br>(Invitrogen) | 1:1000 |
| Rb Glial fibrillary<br>acidic protein (GFAP) | Ab278054<br>(Abcam) | 1:500 |  |  |  |

### 2.4 Extracellular electrophysiological recordings

Neural activity was recorded on the Axion Maestro Pro MEA system (Axion BioSystems, Georgia, USA) with an integrated temperature-controlled CO_2_ incubator (temperature 37°C and 5% CO_2_). Data acquisition was done through the AxIS Navigator Software version 3.1.1. Spontaneous neuronal activity was recorded across 5 weeks between 9 DIV and 32 DIV. Spiking data was captured using the AxIS spike detector with an Adaptive threshold crossing. Spikes were defined by a threshold of 7 standard deviations of the internal noise level with a post/pre-spike duration of 2.16/2.84 ms of each individual electrode, and with frequency limits of 200Hz – 3kHz. Spike sorting was not attempted due to high clustering of the neurons on each electrode making it challenging to reliably discern which spikes correspond to individual neurons on the electrode. Furthermore, we were interested in the network wide activity rather than the activity of individual neurons.

### 2.5 Chemogenetic manipulation

To investigate the network response to chemogenetic manipulation, the novel synthetic ligand DCZ (MedChemExpress) was used to activate the DREADDs receptors (Nagai et al., 2020, Bjorkli et al., 2022) to induce synaptic silencing in excitatory neurons (see **Figure 2** for workflow). In summary, MEA plates were incubated for 15 minutes in the Maestro Pro chamber to allow the activity to stabilize before commencing the recording. Then, baseline activity was recorded for 20 minutes to capture the spontaneous activity of the networks before either PBS or DCZ was added. Afterwards, either PBS (vehicle) or DCZ diluted in cell media (treatment) was added to 45% media volume in the wells at a final DCZ concentration of 10 μM. Networks were incubated for 1hour and then recorded for 1hour. This 1hour recording was divided into 3 phases of 20 minutes recordings denoted as 1^st^ Treatment phase, 2^nd^ Treatment phase and 3^rd^ Treatment phase. The recording was continuous, and the division was done offline during the analysis. After treatment, 3 × 50% media changes were performed to wash out DCZ in the inhibited networks, and 3 × 50% media changes done in the PBS treated networks. To keep all conditions similar, a full media change was carried out on the Control (CTRL) networks. Networks were recorded after washout at 12 hours and 24 hours (see **Table 2** for overview of networks recordings and analysis done). We looked at a total of 23 networks across repeated experiments, and 17 networks from the same experiment are presented here in the main results. Six networks were excluded from the main results due to missing data points.

**Figure 2.**
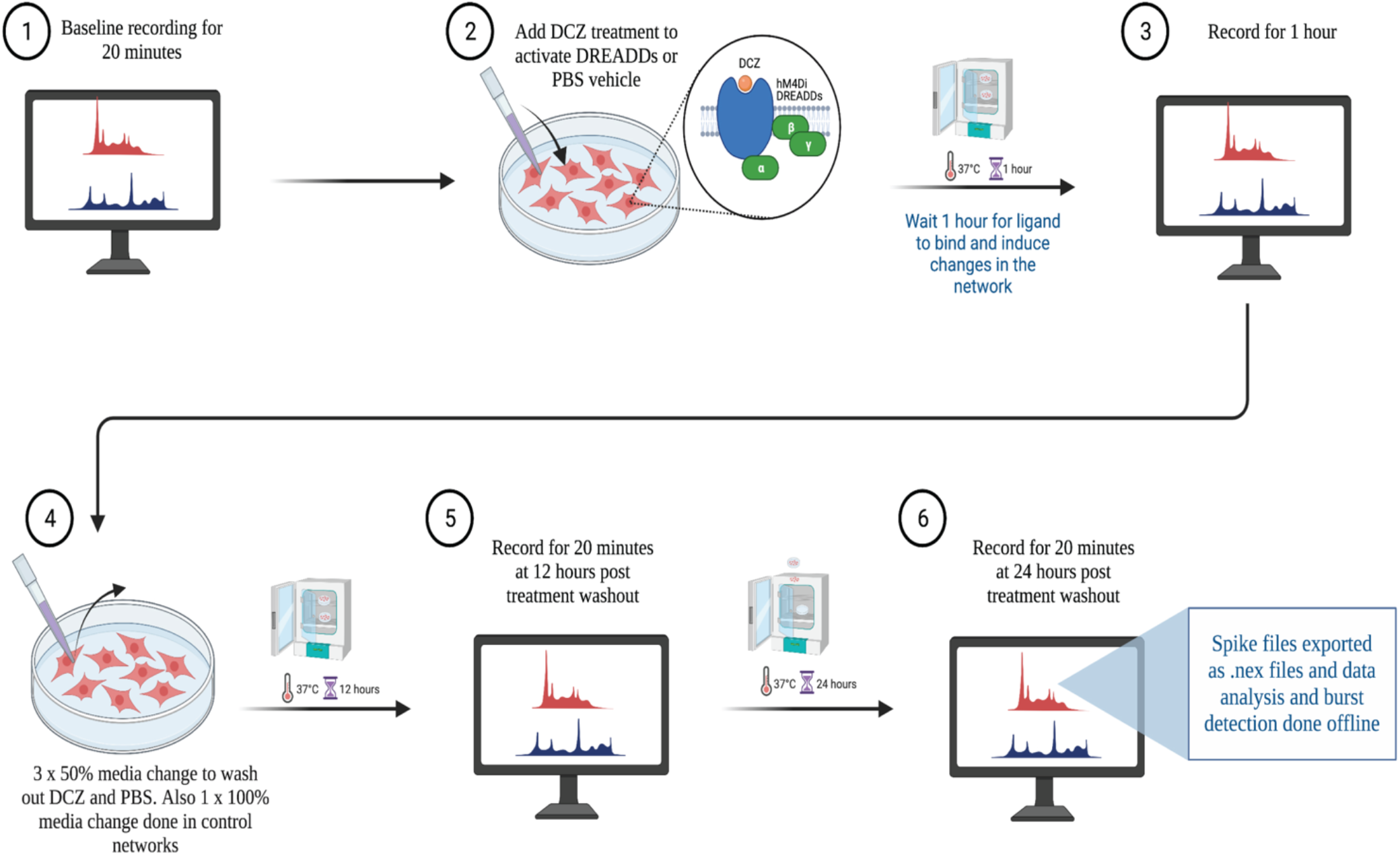
Workflow of electrophysiological recordings with treatment and wash out steps. Created with BioRender.com

**Table 2.** Overview of networks analyzed, conditions, treatments received, and recordings done.

| Neuronal networks analyzed | Conditions | Protocol | Electrophysiology recording |
| --- | --- | --- | --- |
| Control x 7 networks<br>(6 included in main results) | No DREADDs, No DCZ<br>treatment, No PBS vehicle | - | Baseline |
| Control x 7 networks<br>(5 included in main results) | DREADDs + PBS vehicle | 10% PBS in cell media | Baseline<br>During PBS vehicle<br>Recovery at 12hrs<br>Recovery at 24hrs |
| Treatment x 9 networks<br>(6 included in main results) | DREADDs + DCZ treatment | 10uM DCZ diluted in cell<br>media | Baseline<br>During DCZ treatment<br>Recovery at 12hrs<br>Recovery at 24hrs |

**Table 3.**
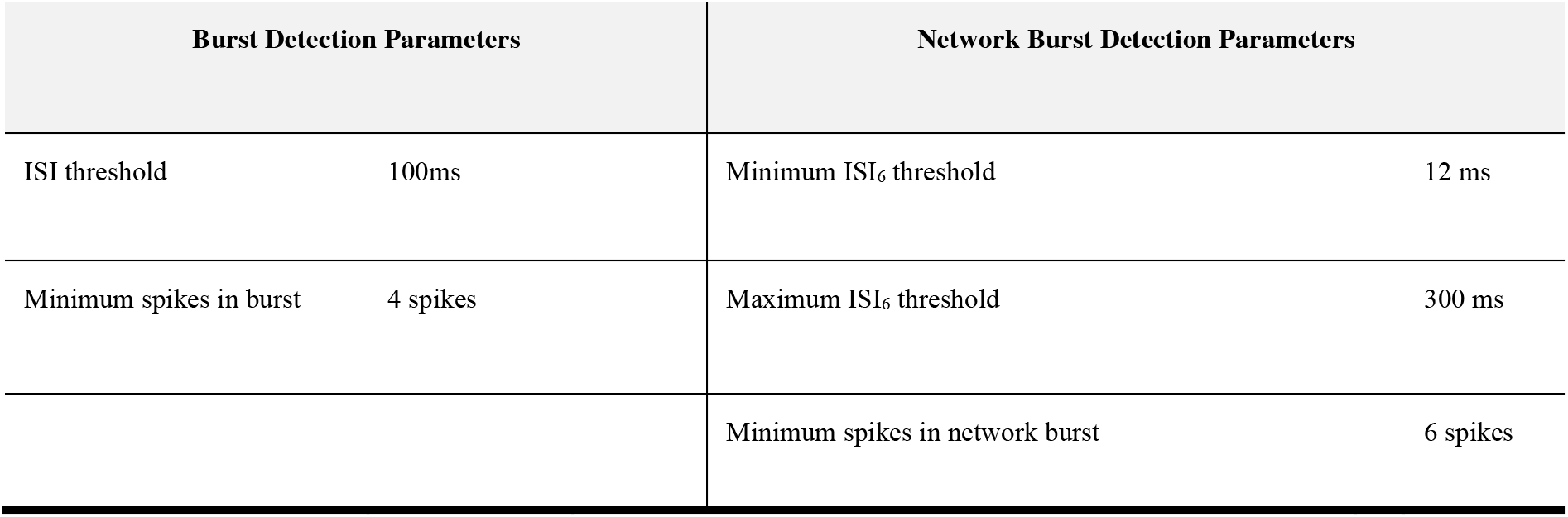
Burst and Network Burst detection parameters on the cumulative spike train over all electrodes.

### 2.6 Network dynamics analysis and network burst detection

The recording spike frequency was computed using the equation: 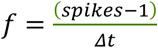, where *spikes* are the total number of spikes for a recording channel and *Δt* is the time difference between the first and last spike included in *spikes*.

For shorter windows we define the instantaneous spike frequency of the window, *f_window_*, as 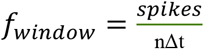.

Here, *spikes* are the spikes in the given window, *n* is the number of active electrodes in the recording, and Δt is the width of the window of interest. The instantaneous spike frequency was computed using a moving window of 1 second with a step size of 0.1 second, resulting in an overlap of windows for instantaneous measures.

Bursts were defined as sequences of at least four spikes with an Inter Spike Interval (ISI) lower than a threshold of 100 ms for all electrodes. The ISI was defined as the quiescent period between two consecutive spikes. Network bursts were defined as the collective sequences of synchronized bursts within an automatically detected ISI threshold for each well at every recording time (Bakkum et al., 2013). First, the ISI between six consecutive spikes (ISI_6_) on the flattened spike train were binned on a logarithmic scale, and the peaks of the binned histograms were detected. The thresholds were centered between these two peaks on a logarithmic scale and limited to the range between at minimum 12 ms and at maximum 300 ms (Gandolfo et al., 2010, Obien et al., 2015). A network burst was detected for spikes where the interval between six consecutive spikes was below the found threshold. The Inter Burst Interval (IBI) was detected as the quiescent period between two bursts or two network bursts (NIBI). Burst analyses were also performed to identify the number of spikes in each network burst (spikes in network burst) and the count of the number of network bursts generated with the number of spikes (number of occurrence). The Burstiness Index of a recording was defined as the amount of activity contained in the 15% most active windows of the computed instantaneous spike frequencies and provides an indication of synchronized neuronal participation in global network bursts (Wagenaar et al., 2005).

The Coherence Index was calculated as the standard deviation divided by the mean of the instantaneous spike frequencies. A high Coherence Index indicated more activity was contained in co-occurring bursts on multiple electrodes. Each parameter of all recording groups was assessed for normality using the Shapiro-Wilk test. Comparisons between groups were evaluated using the Welch’s t-test or the Conover test in the case of normality and non-normality, respectively. Both tests were corrected with Bonferroni corrections for multiple comparisons. Statistical significance was determined if the p-value falls below the significance level (p < 0.05).

## 3 Results

### 3.1 AAV2/1 Gi-DREADD is expressed exclusively in CaMKlla positive neurons

AAV mediated-DREADDs expression was confirmed with immunolabeling to amplify mCherry expression in target CaMKlla positive neurons (**Figure 3A**). Neither inhibitory neurons (GAD65/67) (**Figure 3B**), nor astrocytes (GFAP) (**Figure 3C**) showed co-labelling with mCherry. This confirmed that there was cell specific expression of the AAV-DREADDs. Furthermore, networks at 14 DIV positively expressed GABA (**Figure 4A**), GABA B receptors (**Figure 4B**) and NMDA receptors (**Figure 4C**) confirming network capacity for excitatory and inhibitory signaling at this age.

**Figure 3.**
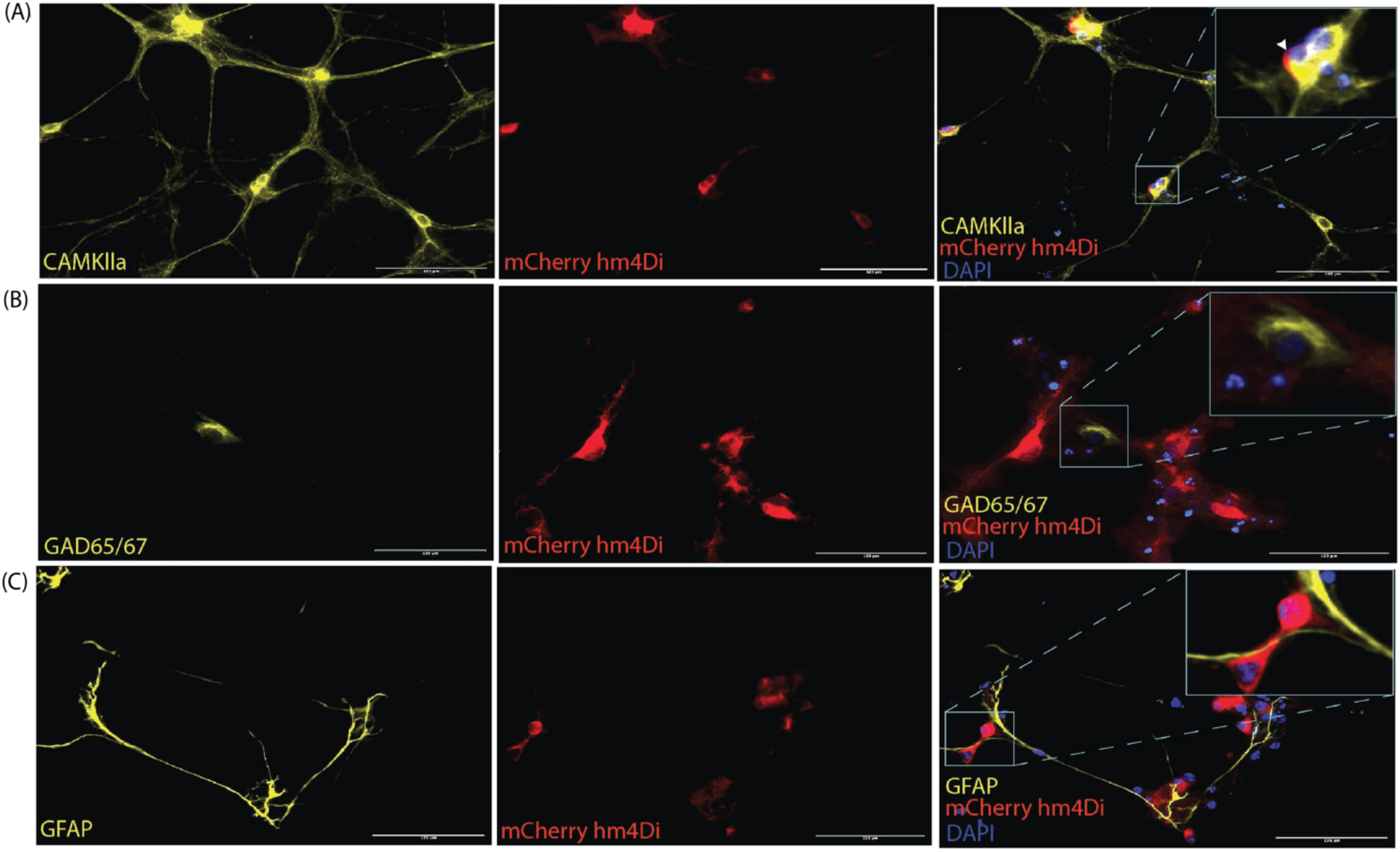
AAV2/1 hM4Di DREADDs expression in neurons in vitro. **(A)** The mCherry antibody was used to enhance the fluorescent of the hM4Di receptors, which were positively colocalized with the somata of CaMKlla positive neurons. **(B)** GAD65/67 expression indicated the presence of inhibitory neurons and showed no soma colocalization mCherry hM4Di expression. **(C)** GFAP antibody was used to label astrocytes in the culture which also showed no soma colocalization with mCherry hM4Di expression. Scale bar = 125μm.

**Figure 4.**
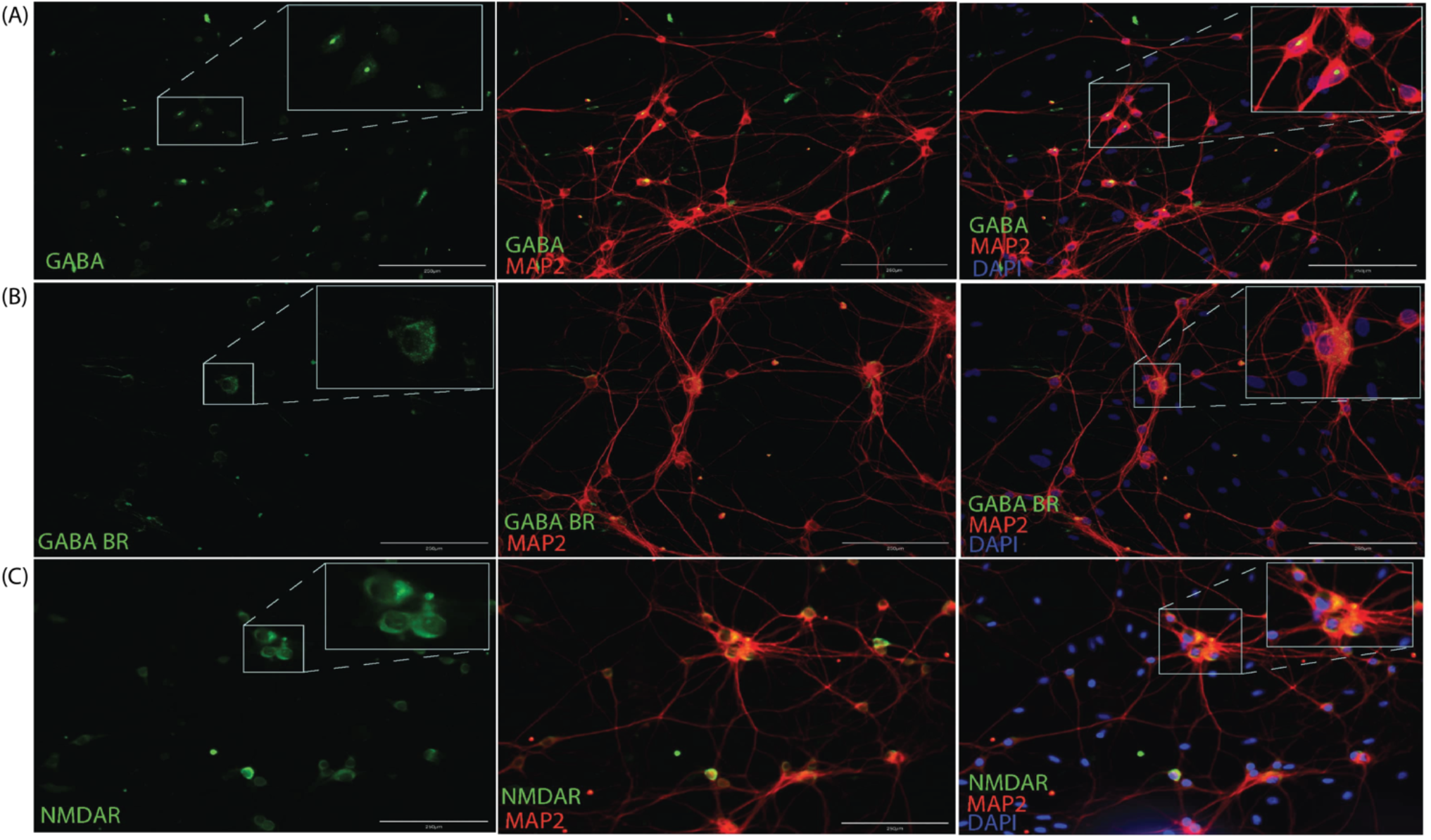
Immunocytochemistry for GABA **(A)**, GABAB receptors **(B)** and NMDA receptors **(C)** along with MAP2 neuronal cytoskeletal marker at 14 DIV. Scale bar 200 μm.

### 3.2 Spontaneous activity and burst characteristics at baseline

Spontaneous network activity was recorded at different timepoints during the experiment for the chemogenetically inhibited networks, PBS vehicle networks and Control networks, which did not receive any treatment (hereafter referred to as DCZ networks, PBS networks and CTRL networks respectively). The spontaneous baseline network profile before the addition of either PBS or DCZ **(Step 1, in Figure 2)** captured across 5 weeks is presented in **Figure 5**. The networks in each condition showed some variations in their activity and bursting characteristics between each recording from 9 DIV to 32 DIV, nonetheless, the mean spontaneous network activity of all networks followed a typical trajectory of development, with increasingly more bursts as the networks reached maturity, according to previous work (Kamioka et al., 1996, Wagenaar et al., 2006). The CTRL networks exhibited more robust electrophysiological activity across several of the parameters, especially in the mean firing rate from 21 DIV onwards when compared to the other networks **(Figure 5 A).** Nonetheless, all networks had a trend of increasing mean firing rate between 9 DIV and 28 DIV with a decrease at 32 DIV **(Figure 5A)**, and an opposite trend in the ISI, which decreased over time until 28DIV, then increased again by 32 DIV **(Figure 5B)**. All networks exhibited bursting activity at 9 DIV and continued to exhibit varying degrees of bursting throughout network lifetime.

**Figure 5.**
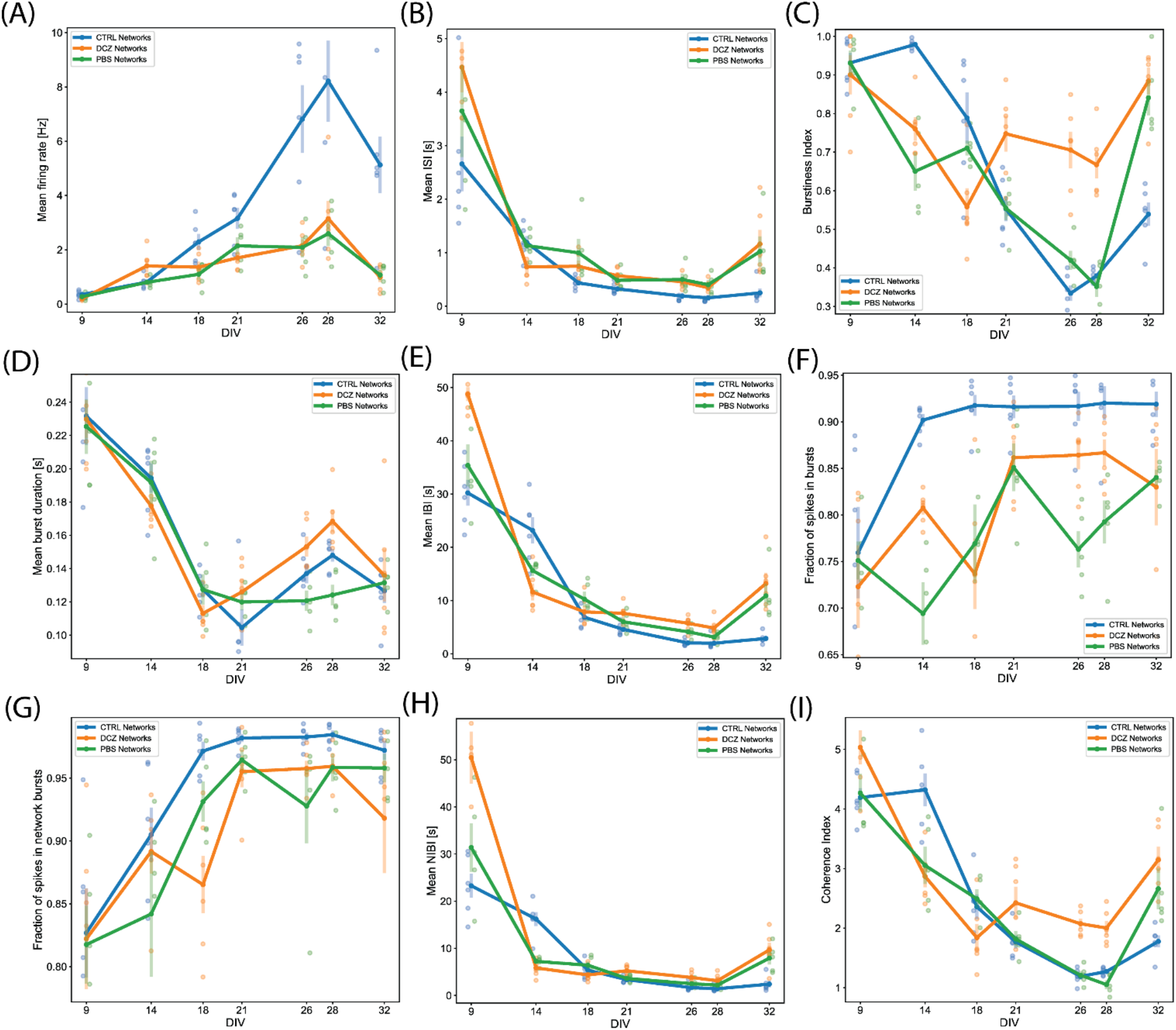
Activity and burst composition at baseline across 5 weeks of recording. Each plot presents the mean activity for all the networks in each condition (DCZ treated (n=6), PBS vehicle (n=5) or CTRL (n=6)). Network behaviour for each condition is described in terms of Mean firing rate **(A)**, Mean interspike intervals (ISI) **(B)**, Burstiness index **(C)**, Mean burst duration **(D)**, Mean interburst intervals (IBI) **(E)**, Fraction of spikes in bursts **(F)**, Fraction of spikes in network bursts **(G)**, Mean network IBI **(H)** and Coherence index **(I)**. The solid lines with solid circles plot the mean values for all networks in one group, the shaded bars show the standard error of the mean, and the shaded circles show the individual data points (the mean activity obtained from each network in each group).

We found that the mean burstiness steadily decreased between 14 and 26 DIV for CTRL networks and between 9 and 28 DIV for PBS networks **(Figure 5C)**. From then onwards, until 32 DIV, both PBS and CTRL networks increased drastically in burstiness. Interestingly, while the DCZ networks also exhibited a decrease in burstiness between 9 and 18 DIV, these networks had a significantly higher burstiness at 21 DIV when compared to PBS (p < 0.02) and CTRL (p < 0.02) networks, and at 28 DIV compared to PBS (p < 0.0006) and CTRL (p < 0.002) networks **(Figure 5C)**. Furthermore, the mean burst duration for all networks across the 3 conditions decreased similarly between 9 and 18 DIV, after which point the DCZ networks started to display increasingly longer bursts, which was significant at 28 DIV when compared to PBS networks (p < 0.003), but not CTRL networks (p > 0.05) **(Figure 5D)**. The CTRL networks also displayed increasingly longer bursts during this time, while PBS networks maintained a stable burst duration between 18 and 32 DIV **(Figure 5D).** All networks maintained a similar trend in mean IBI and mean NIBI, with both decreased steadily between 9 and 28 DIV, with a slight increase at 32 DIV for both DCZ and CTRL networks **(Figure 5E, H)**.

We also noticed that there was a lot of variation between day-to-day recordings in the PBS and DCZ networks for both fraction of spikes in bursts **(Figure 5F)** and fraction of spikes in network bursts **(Figure 5G)**. The CTRL networks however, maintained a very constant burst composition with > 90% spikes occurring in both isolated bursts **(Figure 5F)** and network bursts **(Figure 5G)** from 14 DIV onwards. However, when we looked at network synchrony, which was measured by the coherence index, we noticed that after 18 DIV there was an overall increase in synchrony in DCZ networks at baseline, with a slight decrease between 21 and 28 DIV. Both PBS and CTRL networks exhibited decreased synchrony, with PBS networks decreasing between 9 and 28 DIV and CTRL networks between 14 and 28 DIV **(Figure 5I)**, even though both had > 90% spikes occurring in network bursts from 18 DIV onwards **(Figure 5G)**. The increase observed in DCZ networks at 21 DIV did not differ significantly when compared to the other networks, but there were significant changes at 26 DIV compared to CTRL (p < 0.0007) and PBS (p < 0.002) networks, and at 28 DIV compared to CTRL networks (p < 0.004) and PBS networks (p < 0.0004). This increase in synchrony in DCZ networks from 18 DIV also corresponded to the observed increase in burstiness and burst durations at the same timepoint **(Figure 5C, D)**. At 32 DIV, all networks including PBS and CTRL networks showed an increase in synchrony **(Figure 5I)**, with only a significant difference between DCZ and CTRL networks (p < 0.003).

### 3.3 Analysis of network response and network recovery due to selective inhibition

To identify the changes in activity in the neural networks, we compared spontaneous baseline activity with activity during either DCZ treatment or PBS vehicle, as well as the network activity after DCZ removal at different time intervals. Hereafter, we refer to the recordings during DCZ treatment or PBS vehicle as ‘response’. In these results, we have only included the analysis of the recordings done at 12 hours and 24 hours post washout as we were interested in the network’s recovery over a longer timeframe after perturbation. These recordings will be subsequently referred to as ‘recovery’. The response activity was analyzed in 3 phases of 20 minutes recordings – 1^st^ phase, 2^nd^ phase, and 3^rd^ phase – to better characterize dynamic network changes. The baseline activity and inhibited activity of one DCZ treated network are shown as the recording trace generated from 64 channels on the MEA (**Figure 6A**). Prior to DCZ application, the spontaneous firing rate at baseline was stable for the entirety of the recording, observed as regular spikes and a high occurrence of bursts containing < 10 spikes per bursts (**Figure 6A, first panel labeled ‘Baseline’**; **Figure 6B, C)**. As expected, the application of DCZ caused a decrease in network activity and ablation of networks bursts, which was captured during the 1^st^ phase response (**Figure 6A, second panel labeled ‘Treatment 1^st^ phase’**). The network started exhibiting intermittent spikes and isolated bursts that gradually increased as the recording progressed (**Figure 6A, third and fourth panels labeled ‘Treatment 2^nd^ phase’ and ‘Treatment 3^rd^ phase’**), indicating that network activity recovered in the presence of DCZ. We also noticed that during the 1^st^ phase response, the DCZ networks exhibited very low occurrences of bursts (< 2 occurrences of bursts at any timepoint during the recording period), and the occasional burst had up to 150 spikes per bursts for individual bursts (**Figure 6B**) and up to 800 spikes per bursts for network bursts (**Figure 6C**). There was also an increase in the number of burst occurrences for the 2^nd^ and 3^rd^ phase responses for both individual bursts and network bursts for the DCZ networks, exceeding 600 occurrences of bursts with < 10 spikes in bursts for the 3^rd^ phase response (**Figure 6B**) and up to 200 occurrences of bursts with < 10 spikes in network bursts **(Figure 6C)**. The PBS networks depicted here maintained some bursting activity during the 1^st^ phase response, though there were lower occurrences of bursts and fewer spikes in both individual bursts and network bursts when compared to the DCZ networks (**Figure 6B, C**). There was, however, a gradual increase in the number of spikes in bursts at the 2^nd^ and 3^rd^ phase response for both individual bursts and network bursts **(Figure 6B, C)**. The PBS networks also maintained a trend similar to DCZ networks where the most occurrences of bursts had < 10 spikes, and there were some bursts with up to 100 spikes per burst by the 3^rd^ phase response for both individual bursts and network bursts. Unlike the inhibited networks though, which had up to 1000 spikes per network burst by the 2^nd^ phase response, PBS networks did not exceed 100 spikes in bursts or network bursts (**Figure 6B, C**).

**Figure 6.**
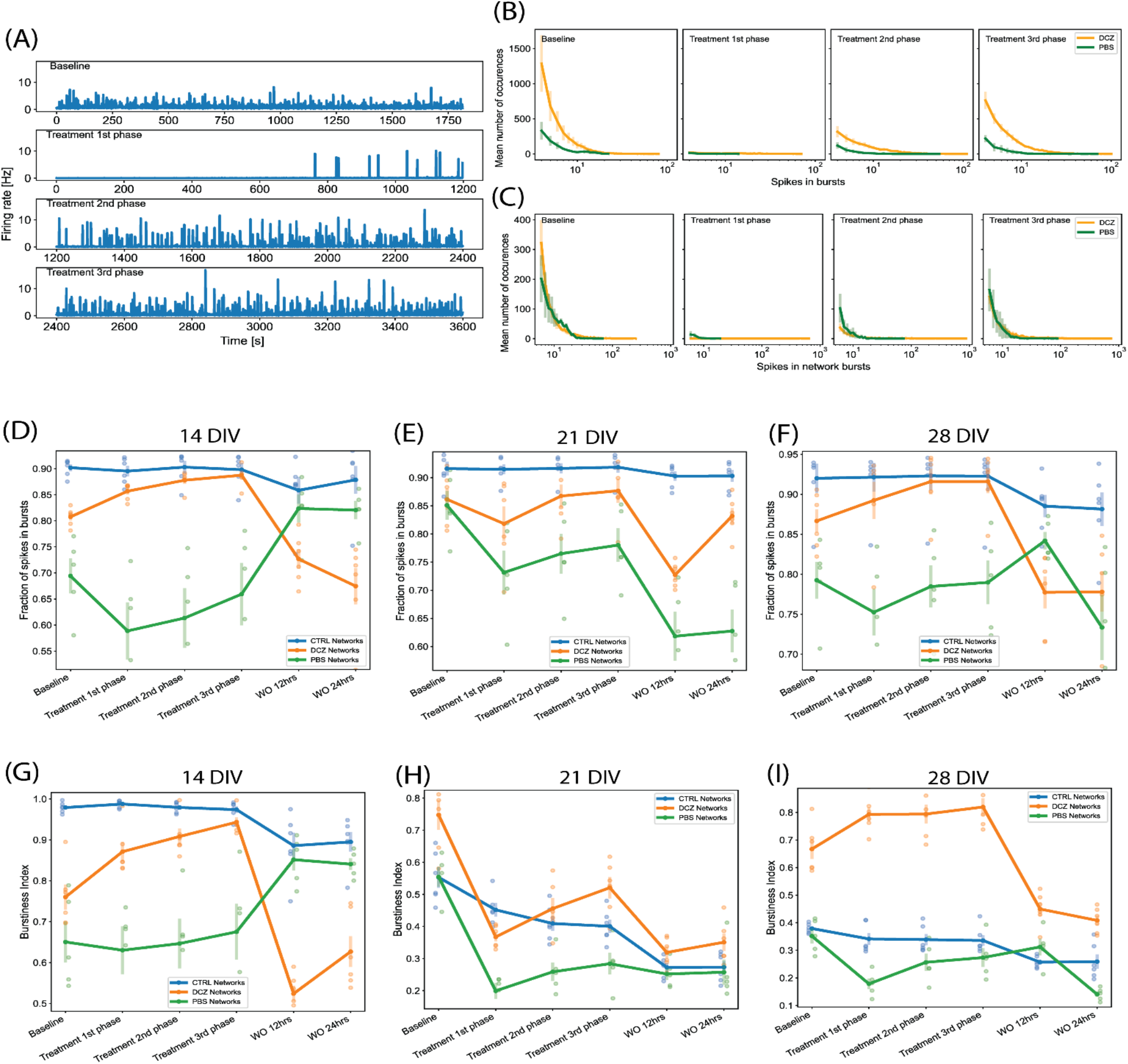
Neural network activity at baseline and in response to DREADDs-mediated inhibition of excitatory synaptic transmission. Each panel in **(A)** show a trace generated from 64 recording channels of spontaneous activity of one DCZ treated network at 28DIV. The first panel shows the 20 minutes recording of the spontaneous firing rate at baseline, the second, third and fourth panels show the 1^st^, 2^nd^, and 3^rd^ phases of 20 minutes recordings of spontaneous activity during DCZ treatment. The x-axis denotes time in seconds and the y-axis denotes firing rate in Hz. **(B)** The spikes in bursts and **(C)** network bursts for the baseline, 1^st^, 2^nd^, and 3^rd^ phase recordings are shown for sample DCZ treated networks (n=3) and for sample PBS vehicle networks (n=2). The x-axis denotes the number of spikes, and the y-axis denotes the number of burst occurrences of a given number of spikes. **(D-F)** Plots of the fraction of spikes in bursts across baseline, treatment and recovery recordings at 14 DIV, 21 DIV and 28 DIV for all network groups (n=6 for CTRL and DCZ, and n=5 for PBS). The x-axis denotes the recording condition, and the y-axis denotes the percentage of spikes located in bursts. **(G-I)** Plots depicting burstiness of each network group across the baseline, treatment and recovery recordings at 14 DIV, 21 DIV and 28 DIV. The x-axis denotes the recording condition, and the y-axis denotes the burstiness index as the fraction of activity in the 15% most active time windows. The solid lines and solid circles plot the mean values for all networks in one group, the shaded bars show the standard error of the mean, and the shaded circles show the individual data points.

We performed further analyses to look at both the fraction of spikes in bursts and the burstiness index for the networks at baseline, during response, and during recovery on the days that they were manipulated (14 DIV, 21 DIV and 28 DIV). These results revealed that the CTRL networks maintained their characteristic of having > 90% of spikes located in bursts across all the recordings (baseline, response, and recovery) at 14, 21 and 28 DIV (**Figure 6D – F**). There were no significant changes in the fraction of spikes in bursts for CTRL networks at recovery. We noticed that there was a decrease in the fraction of spikes in bursts between baseline and 1^st^ phase response across all the days for the PBS networks, and a slight increase during the 1hour response recording (**Figure 6D – F**), however these changes were not found to be significantly different from baseline (p > 0.05). At 14 DIV, the PBS networks had a very quick recovery at 12 hours, exhibiting > 80% of spikes in bursts which was maintained for at least 24 hours. However, recovery at 12 hours appeared impaired at 21 DIV, at which time point the PBS networks decreased significantly below baseline in the fraction of spikes in bursts (p < 0,05) (**Figure 6E**). Interestingly, although DCZ networks had a nonsignificant decrease in the fraction of spikes between baseline and the 1^st^ phase response at 21 DIV, these networks stably maintained > 80% of spikes in bursts between baseline and during the 1hour response recording for all 3 days (**Figure 6D – F**). As expected, there was a significant decrease in the fraction of spikes in bursts after DCZ washout at 12 hours compared to baseline across the 3 perturbation days. This change, however, was only significant at 21 DIV (p < 0.005) and 28 DIV (p < 0.006) (**Figure 6D – F**). While the CTRL networks maintained a high bursting profile at 14 DIV across all the recordings (**Figure 6G**), this steadily decreased until burstiness had diminished significantly by 28 DIV when compared to DCZ networks. PBS networks also had lower burstiness profiles across all the recording sessions at 28 DIV where we saw a distinct difference in burstiness at 2^nd^ and 3^rd^ phase responses compared to DCZ networks (p < 0.00005; p < 0.00003 respectively). The DCZ networks maintained a high burstiness especially noticeable during the 1hour response recording at 14 and 28 DIV **(Figure 6G – I)**. However, at 21 DIV, there was a significant decrease in burstiness between baseline and the 1^st^ phase response (p < 0.00001), and although there was a significant increase between 1^st^ and 3^rd^ phase response (p < 0.006), this was still significantly lower than baseline (p < 0.05). In addition, as can be observed in (**Figure 6D – F**), across all the perturbation days burstiness decreased to significant levels after washout at 12 hours recovery when compared to baseline at 14 DIV (p < 0.002), 21 DIV (p < 0.0000005) and 28 DIV (p < 0.005). Activity in the DCZ networks did not recover to baseline levels within 24 hours (**Figure 6G – I**).

We also found that during response at 14 DIV, the PBS and DCZ networks had overall shorter mean burst duration, and shorter mean IBI than the CTRL networks (**Figure 7A, B**). These differences were found to be significant when comparing DCZ with CTRL networks at 1^st^ (p < 0.04), 2^nd^ (p < 0.04) and 3^rd^ (p < 0.03) phase responses, and PBS and CTRL networks only at 2^nd^ (p < 0.005) and 3^rd^ (p < 0.002) phase responses. There were no significant differences in the responses between DCZ and PBS networks. There was also a decrease in both mean burst duration and mean IBI for CTRL networks at 24 hours recovery, while both DCZ and PBS networks increased in both parameters **(Figure 7A, B)**. At 28 DIV, consistent with what was seen with the burstiness index in **Figure 6I**, the DCZ networks had an overall steady increase in mean burst duration during the 1hour response recording, with correspondingly longer intervals between each burst (**Figure 7 E – F**). DCZ networks also had a decrease in both mean burst duration and mean IBI between 3^rd^ phase response and 12 hours recovery, with a slight increase in mean IBI at 24 hours recovery (**Figure 7E, F**). Interestingly though, there was variability in the responses across the networks, especially observed at 14 and 21 DIV (**Figure 7A – D**). Both days showed an increase in mean burst duration at 12 hours recovery for all networks, but this was sustained until 24 hours only at 14 DIV (**Figure 7A, C**). Similarly, for both mean burst duration and mean IBI at 21 DIV, there were no significant differences in the response between any of the networks across the recordings, though there was an overall decrease in the CTRL networks compared to what was observed at 14 DIV.

**Figure 7.**
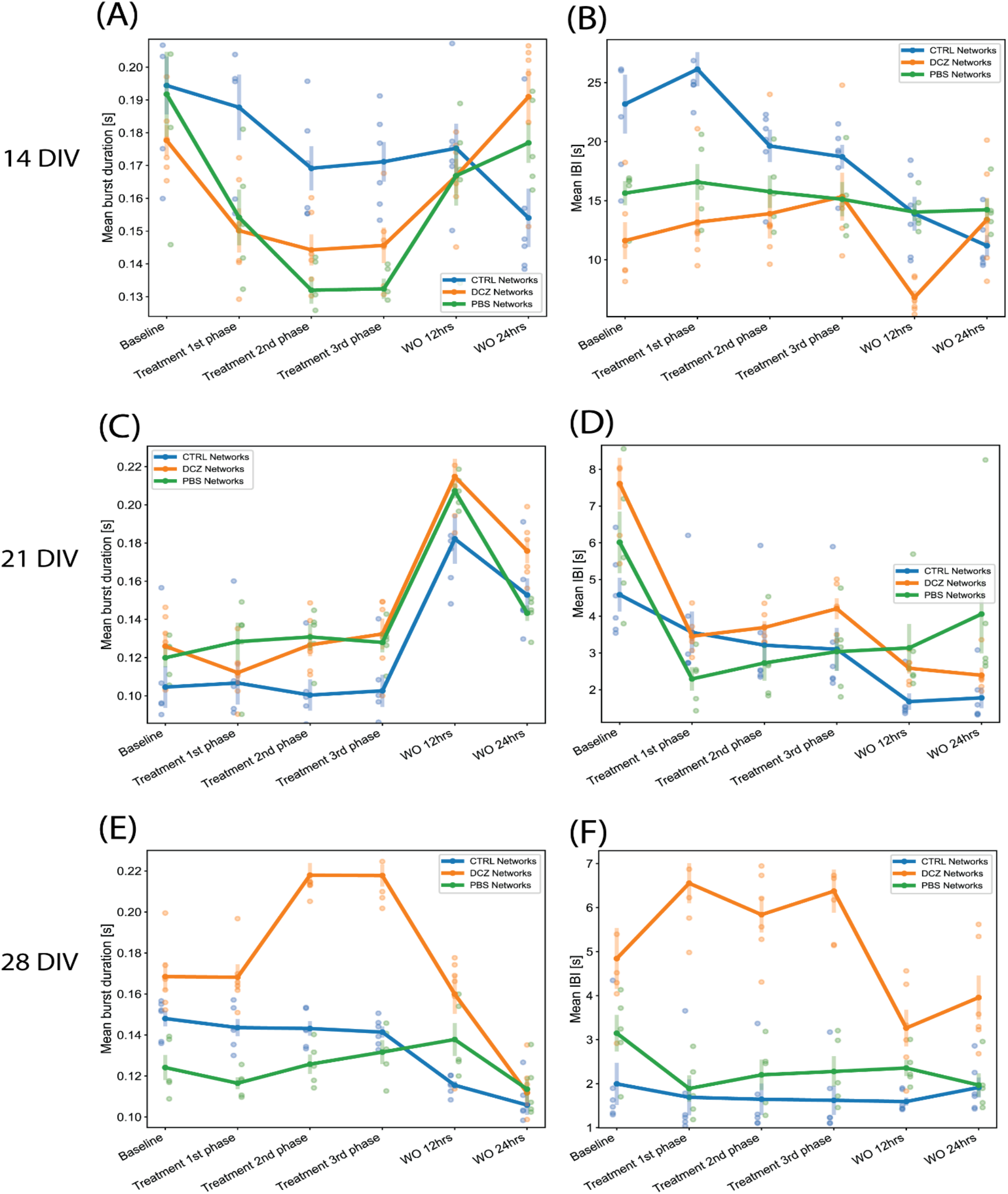
Neural network mean burst duration and mean interburst intervals. (**A, C, E**) Plots showing the mean burst duration and (**B, D, F**) showing the mean IBIs for each network group (n=6 for CTRL and DCZ, and n=5 for PBS) across the baseline, treatment and recovery recordings at 14 DIV, 21 DIV and 28 DIV. The solid lines and solid circles plot the mean values for all networks in one group, the shaded bars show the standard error of the mean, and the shaded circles show the individual data points.

### 3.4 Analysis of network bursts and synchrony

Since we observed that the increase in bursting activity in DCZ networks during response seemed to be a result of selective silencing, we wanted to investigate how synchronous the networks were across the different recording phases in comparison to the PBS and CTRL networks. Again, we observed that the CTRL networks exhibited between 90% and 98% of spikes consistently in network bursts across the different recording sessions and for all perturbation days (**Figure 8A, C, E**). However, there was notable variability in the coherence index between the networks at 14 and 21 DIV, with CTRL networks having highest values across the response phases at 14 DIV **(Figure 8B)**. However, synchrony gradually decreased for both CTRL and PBS networks until 28 DIV, but increased for DCZ networks (**Figure 8 B, D, F**). Though the fraction of spikes in network bursts for PBS networks decreased between the baseline recording and the 1^st^ phase response on all days, this was only found to be significant at 21 DIV (p < 0.02) (**Figure 8A, C, E**). The PBS networks also maintained lower synchrony than the DCZ networks during response across all days (**Figure 8B, D, F**). Additionally, for all the perturbation days, the DCZ networks maintained > 90% spikes in network bursts during the 1hour response recording but they did not fully recover to baseline after the media changes at 12 or 24 hours (**Figure 8A, C, E**). Similarly, the DCZ networks also had sustained synchrony during the 1hour response recording, but reduced synchrony at 12- and 24-hours recovery for all 3 perturbation days **(Figure 8B, D, F).** Overall, these results indicate that the inhibited networks steadily began developing more synchronous activity after the first perturbation session at 14 DIV but failed to recover baseline dynamics within 24 hours after the perturbation.

**Figure 8.**
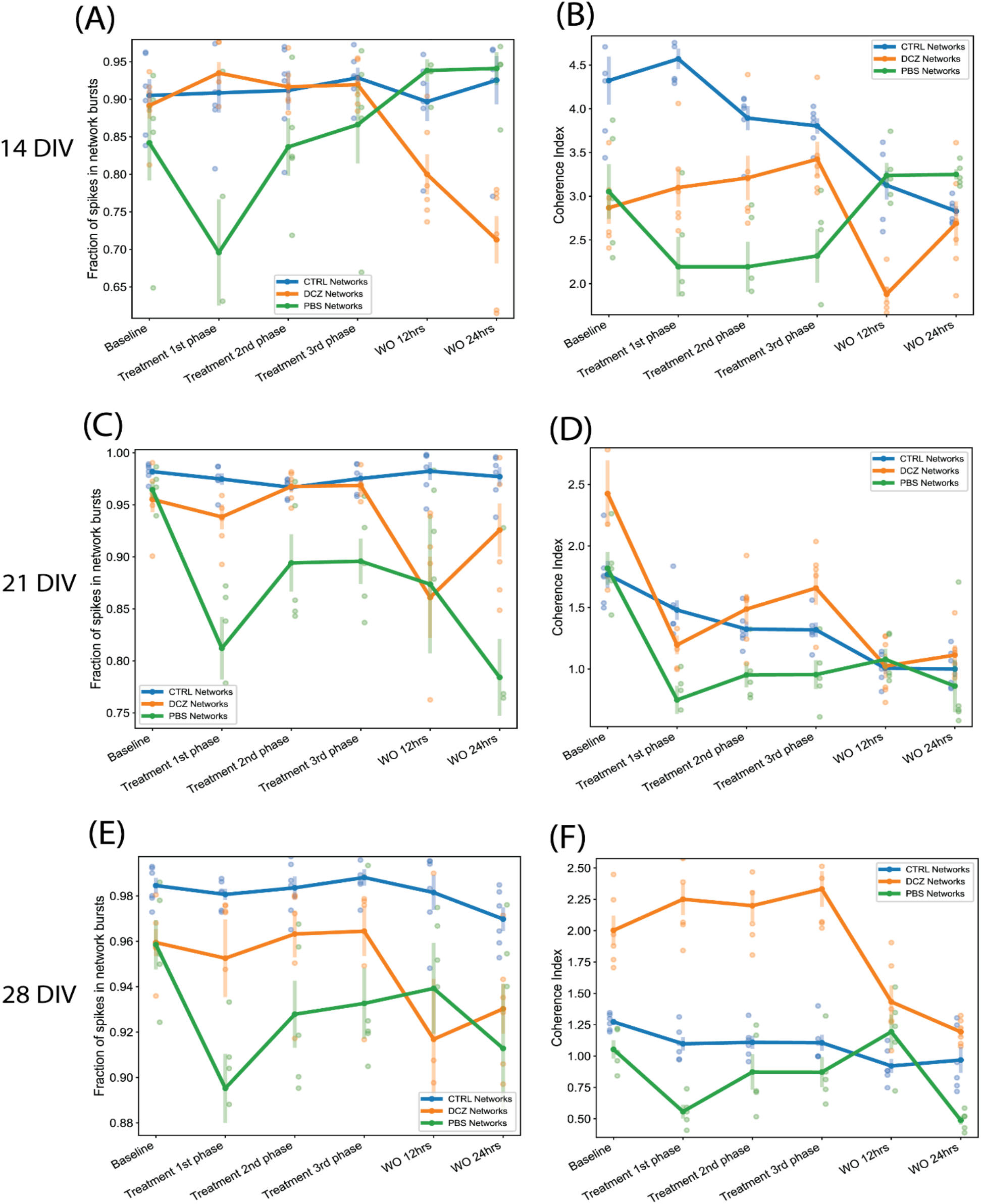
The fraction of spikes in network bursts and the measure of network synchrony across the baseline, treatment and recovery recordings at 14 DIV, 21 DIV and 28 DIV. **(A, C, E)** Plots showing the fraction of spike in network bursts. The x-axis denotes the recording condition, and the y-axis denotes the percentage of spikes in network bursts. **(B, D, F)** Plots showing the coherence index (y-axis) of each network group across each recording condition (x-axis). The solid lines and solid circles plot the mean values for all networks in one group (n=6 for CTRL and DCZ, and n=5 for PBS), the shaded bars show the standard error of the mean, and the shaded circles show the individual data points.

## 4 Discussion

Over the last decades, an increasing amount of research is conducted to answer questions related to in vitro neural network development, E/I interaction, and observed spontaneous dynamic network properties in the absence of external stimuli (Latham et al., 2000). Cortical neurons in vitro tend to form densely connected networks by 7 DIV, as observed in this study (**Figure 1)**, and by 14 DIV, the neurons had formed distinct structural organization with prominent axon fasciculation, and dendritic connections across the entire network, as well as mature excitatory and inhibitory receptors as seen in **Figure 3**. A recent study has shown that functional interactions between maturing excitatory and inhibitory synapses result in dynamic spiking activity and the emergence of network bursts (Teppola et al., 2019). Increasing either excitation or inhibition can therefore be expected to result in aberrant bursting dynamics in neural networks, thus we set out to investigate how bursting dynamics are affected and how neural networks recover when excitatory synaptic transmission is transiently inhibited. To do this, we took advantage of the unique opportunity that DREADDs provide to selectively target excitatory activity, and after transducing the networks with AAV 2/1 hM4Di CaMKlla-DREADDs, we proceeded to activate the DREADDs with DCZ at 14 DIV, 21 DIV and 28 DIV. Our primary findings are: (1) Inhibition of excitatory synaptic transmission resulted in an increase in network burstiness by 28 DIV; (2) Inhibited networks recovered activity in the presence of DCZ indicating rapid homeostatic response to network silencing; (3) By 28 DIV, inhibited networks exhibited higher synchrony and burstiness during and following selective inhibition contrary to PBS and CTRL networks that had diminished levels.

Network activity and bursting dynamics are inherently unique to each network in vitro, nonetheless, in our study all networks exhibited some degree of network bursting activity by 9 DIV. Early network bursts are significant for network development and maturity and are deemed to be physiologically relevant for neural information processing and synaptic plasticity (Lisman, 1997). In developing networks, bursts act as more reliable determinants of neurotransmitter release than single spikes (Lisman, 1997, Delattre et al., 2015), thus synaptic efficacy and facilitation rely on network bursts to increase the probability of postsynaptic response to presynaptic inputs. While others (Marom and Shahaf, 2002, Wagenaar et al., 2006, Chiappalone et al., 2006, Bisio et al., 2014) reported increase in network bursts towards more mature stages in vitro (21 – 28 DIV), our networks showed a propensity towards high, regular bursting activity – as can be seen in **Figure 5 F, G** where over 70% of spikes occurred in bursts and network bursts – for all networks from as early as 9 DIV. Due to their early appearance, these bursts appeared to be akin to ‘superbursts’ typically observed at earlier development, before the network establishes more mature neuronal phenotypes and before GABA receptors mature (Stephens et al., 2012), and may be driven by the early evolution of the network morphology (Kim and Lee, 2022).

Evolving network morphology may play a significant role in the electrophysiological dynamics of the networks throughout development. Neural networks develop and mature through a bottom-up process of self-organization which can be observed everywhere in nature, from the microscopic to the macroscopic level (Turing, 1990, Kondo and Asal, 1995, Arango-Restrepo et al., 2021). The process of self-organization involves the dynamic interaction between constituent elements of a system and implies that there is a reciprocal relationship between structural organization and function (Karsenti, 2008). In physical and biological systems, self-organization is part of emergence, i.e., unpredictable interactions between known constituent elements, and drives morphogenesis (Chialvo, 2010, Dobrescu and Purcarea, 2011). Inherent to the process of self-organization of neural networks is the gradual development of complex hierarchies through local interactions (Karsenti, 2008, Sasai, 2013). Thus, each neural network can be expected to self-organize in a different way. This may explain the observed variability in the baseline activity between each experimental group, as well as between recordings from the same group as shown in **Figure 5.** It is reasonable to assume that each in vitro neural network will have unique structural and functional features, such as dendritic – axonal topological arrangement, cell body clustering and synaptic connections. This will ultimately influence the firing properties and the overall activity dynamics of the network, and in turn, the differences in activity may regulate morphological development (Kater et al., 1988, Kater et al., 1994).

As we observed, the CTRL networks exhibited significantly higher activity especially in the mean firing rate between 18 DIV and 32 DIV and sustained very high percentages of spikes located in either bursts or network bursts across all the recording days when compared to PBS and DCZ networks. In addition, it was also evident that between 9 and 21 DIV, the CTRL networks exhibited significantly higher burstiness than both PBS and DCZ network groups, as well as higher synchrony at 14 DIV than both groups as noted in **Figure 5.** This high activity may be mediated by the topological organization of neurons on the MEAs. There is evidence supporting the idea that mesoscale network architecture and local neuron clustering shape the patterns of persistent local network activity in ongoing network activity (Kaiser and Hilgetag, 2010, Klinshov et al., 2014). Neurons within clusters may receive stronger synaptic inputs, exhibit more intense activity, and contribute more to the initiation, propagation, and maintenance of activity (Okujeni et al., 2017). To add to the complexity, as in vitro networks mature, the firing rate profiles can fluctuate greatly among different sites in the network, giving widely varying electrophysiological profiles for the same network (van Pelt et al., 2004, Habets et al., 1987). We can still, however, confidently draw comparisons between neural networks particularly because intrinsic temporal developmental programs (for example E/I synaptic development) govern the formation of all networks (Ben-Ari, 2001), so they all reliably exhibit consistent patterns of age dependent bursting behaviour, making bursting a reliable measure of network development and maturity, as well as network function and potential dysfunction.

It is hardly surprising that the developmental profile of total network firing and bursting activity vary from recording to recording between the networks. It should be noted that because neural activity is spontaneous and unpredictable, electrophysiological data obtained within narrow study timeframes for example < 28 DIV (Weir et al., 2015, Passaro et al., 2021), and recording time frames for example < 10 minutes recordings (Jimbo et al., 1999, Eytan et al., 2003, Passaro et al., 2021) may present more uniform behaviour and not adequately reflect dynamic network changes. As such, variability in network activity may currently be underreported in the relevant literature when networks are not monitored over longer recording timeframes. This highlights a necessity for more studies that show variability in network electrophysiological profiles over time. In our study we monitored network activity from early development, until 32 DIV, a time frame widely accepted as a period of network maturity (Wagenaar et al., 2006). In addition, we recorded continuous spontaneous baseline activity for 20 minutes and, response activity for 1hour as opposed to 3-10 minutes recordings often reported in the literature. Our longer recordings make it easier to capture variable profiles in network activity.

Notwithstanding the variability in network activity profiles, the responses of the DCZ networks were consistent and distinct from the CTRL and PBS networks and demonstrate that selective inhibition of excitatory synaptic transmission can modulate long term network dynamics. We found that network burstiness began increasing steadily between the first and second perturbation session in DCZ networks and remained high while the PBS and CTRL networks decreased in burstiness as shown in **Figure 5C**, suggesting that selective inhibition affected the maintenance of endogenous network excitation and inhibition, and affected network bursting. Importantly, both PBS and CTRL networks showed a sustained decrease in baseline burstiness over time, as well as an overall decrease in baseline synchrony. This indicated that while bursting may be the dominant activity profile for these networks, there was still a dynamic balance being maintained between E/I, such that global inhibition may have played a role in desynchronizing the network, which may be a fundamental process in neural network development. According to studies investigating sensory coding, desynchronization in neural networks optimizes information processing and performance (Waschke et al., 2019) and may strongly improve the fidelity with which novel information is encoded (Pachitariu et al., 2015). Increased synchronization is implicated in several neurological disorders including but not limited to epilepsy and Parkinson’s disease, where inhibition becomes severely impaired (Calcagnotto et al., 2005, Harrington et al., 2018). Thus, it follows that the uninhibited networks would mature and develop the appropriate excitatory and inhibitory processes necessary to maintain network activity within a healthy dynamic range and achieve desynchronization in order to optimize network information processing capabilities. The observed decrease in coherence in the DCZ networks between 9 and 18 DIV reflected what was observed in the uninhibited networks as part of the normal process of development. It is possible that inhibition at 14 DIV may have triggered the slow synaptic plasticity process mediated by G-protein coupled signaling systems to, for example, induce long term modification of pre and postsynaptic inhibitory response (Chiu and Weliky, 2001, Rozov et al., 2017, Chiu et al., 2019) Therefore, we conclude that transient external inhibition may trigger the network to decrease endogenous inhibitory mechanisms leading to an overall increase in global activation of the neural network.

While there may be different explanations as to the cause of an increase in synchronization and a decrease in inhibition, the most plausible one may be linked to our experimental set up and methods used. In our study, the activation of hM4Di DREADDs blocks cAMP production (by Gai protein blockade of adenylate cyclase), which results in neurons being unable to detect and respond to extracellular signals. Thus, DREADDs expression and activation on excitatory neurons likely prevents neurons from reliably responding to excitatory post synaptic potentials, thereby causing disruption in activity, and the potential development of inhibitory synapses. It is well documented that excitatory synaptic activity regulates the development and maintenance of inhibitory synapses on excitatory neurons (Lin et al., 2008), and that deprivation of excitatory synaptic activity reduces the density of synaptic GABA receptors, and the number of functional inhibitory synapses in cortical cultures (Kilman et al., 2002) and hippocampal slices (Ramakers et al., 1994, Muramoto et al., 1996, Chub and O’Donovan, 1998). In our study, the consequence of excitatory synaptic inhibition at 14 DIV was a subsequent increase in burstiness and synchrony in DCZ networks at baseline, indicating impaired inhibitory synaptic development and overall, less inhibition in the network.

Although the emerging picture is that E/I synaptic activity is the single most important factor regulating neural network bursting behaviour, our results also indicate that there are intrinsic homeostatic mechanisms at work. This is especially relevant considering the recordings during response and recovery at the different perturbation days for the DCZ networks (**Figure 6 - 8)**. According to the theory of homeostatic plasticity, network activity is stabilized by a negative feedback process where a forceful change in activity is resisted, and the system returns to a tolerated dynamic range (Turrigiano, 1999). The data presented in our study show that inhibited networks were able to recover network bursts during DCZ exposure, supporting several previous studies where networks bursts were maintained in the presence of activity suppressing chemogens (Chub and O’Donovan, 1998, Li et al., 2007, Zeldenrust et al., 2018). The exact mechanism for recovery during chemogenetic manipulation is unknown, however, we posit that several homeostatic factors including alterations in neuromodulator levels and neurotransmitter release (Ramakers et al., 1994, Muramoto et al., 1996, Chub and O’Donovan, 1998) or sensitivity (Turrigiano et al., 1998, Desai et al., 1999) contributed to the network rescuing spontaneous activity.

Furthermore, an increase in burstiness and synchrony during DCZ silencing may indicate that silencing excitatory synaptic transmission may have lowered the spike threshold of excitatory neurons causing neurons to respond more robustly to homeostatic activation, in a manner that reverberates in the network without much inhibitory control. We know from this study and others that in vitro, neurons tend to connect with each other in a modular organization of several clusters connected by both long- and short-range connections (Antonello et al., 2022). Within a network with reduced inhibition, as one module becomes activated whether spontaneously or due to external influence, the activity will quickly spread throughout the network in a positive feedback manner, increasing network synchronization (Huang et al., 2017). Our results also suggest that homeostatic mechanisms might play a role in the recovery of the DCZ networks at 28 DIV as seen with the decrease in burst duration and IBIs (**Figure 7**) as well as burstiness and synchrony (**Figure 8**) between 3^rd^ phase response and 12 hours recovery. We cannot entirely exclude however that such changes may be related to the media changes done in order to wash out DCZ from the networks. Also, though activity recovered in the sense that there was a decrease in burstiness and synchrony, the inhibited networks did not recover to baseline, but rather had drastically lower activity at both 12- and 24-hours recovery as shown in **Figure 6, 7 and 8.** This may indicate that recovery to baseline is a very slow process and takes longer than 24 hours, especially before the networks reach 28 DIV. Since there was an increase in both baseline burstiness and coherence between 28 and 32 DIV for all networks as shown in **Figure 5**, it would be interesting to see whether this would stabilize as the networks get older and remain unperturbed and unstimulated.

Finally, an unexpected observation was a response to PBS vehicle between the baseline and 1^st^ phase responses in PBS networks. PBS is often used as a vehicle in many in vitro and in vivo experiments. Addition of 10% PBS as a vehicle might have affected the concentration of media nutrients and caused a response in the firing activity. On the other hand, the observed effects may be merely due to intrinsic differences in each network in the PBS group. As it relates to the DCZ networks and the variability in the response between baseline and treatment 1^st^ phase especially at 14 and 21 DIV, we cannot rule out that this may be due to where the DREADDs hM4Di are located in the network, and how they get activated. Although several protocols were tested to optimize the concentration of AAV DREADDs and DCZ ligand, we cannot be certain that the same DREADDs on the same neurons, or even on the same part of the network were being activated every time. To our knowledge, this combination using AAV DREADDs, and the novel synthetic ligand DCZ has not been used *in vitro* with dissociated primary neurons, so there are still great possibilities to explore in this area of research.

## 5 Conclusions and future directions

In this study, we investigated the responses of in vitro neural networks to transient selective inhibition of excitatory synaptic transmission, and network recovery from perturbation. We examined characteristics of network bursting dynamics over time, as well as network burstiness and synchrony. We found that while uninhibited networks developed with most of their spikes located in network bursts, inhibited networks overall exhibited more burstiness and synchrony at maturity. The burstiness and synchrony was also maintained during network response recordings, indicating homeostatic mechanisms restoring network activity in the presence of the ligand. The overall increase in burstiness and synchrony after the first perturbation, may be due to a decrease in endogenous inhibitory mechanisms caused by long term inhibitory synaptic modifications. In future studies it will be interesting to monitor the networks in the long term to see how the recovery profile changes with network maturity. As well as investigate the long-term implications of excitatory synaptic silencing on functional connectivity. There might have been some remodeling of synaptic attributes and/or reorganization of the structural network, which would make the network less efficient at information transmission due to the increased synchrony. This hypothesis can be tested further using high density MEAs that offer higher spatial resolution for network connectivity investigation.

## Data availability statement

The raw data that support the findings of this study will be made available by the authors.

## Author Contributions

**Janelle Shari Weir**: Conceptualization, Investigation (Cell experiments, Protocol development and optimization; AAV investigations, Immunocytochemistry and Electrophysiology recordings), Data visualization, Writing – Original draft, Review and Editing. **Nicholas Christiansen:** Methodology, Data analysis (Preprocessing, Scripts, Visualization), Statistical analyses, Writing – Review and Editing. **Axel Sandvig**: Conceptualization, Funding Acquisition, Writing – Review and Editing, Supervision. **Ioanna Sandvig**: Conceptualization, Funding Acquisition; Writing – Review and Editing, Supervision.

## Acknowledgments

The authors would like to thank Prof. Morten Høydal, Department of Circulation and Medical Imaging, NTNU, for facilitating access to the Axion electrophysiology system, and Dr. Rajeevkumar Nair Raveendran, Viral Core Facility, Kavli Institute for Systems Neuroscience, NTNU, for the design and preparation of the AAV DREADDs constructs.

